# Assessing assay absorption artefacts in *in vitro* cell responses to particles

**DOI:** 10.1101/2023.01.16.524328

**Authors:** Seiha Yen, Graeme R. Zosky, Yong Song

## Abstract

In this study, we assessed the issue of coal particles absorbing extracellular proteins and tested the effects of different culture conditions and processing strategies to address this issue. Our data show that there is no effective strategy to solve this problem. We agree with previous reports that cytokine binding experiments should be performed to implement appropriate correction factors in order to calculate the accurate production of secreted cytokines in the supernatant of cell culture experiments. This is an underappreciated issue in many published studies on the comparative potency of particles from different sources.

---

*To the Editor:*

*In vitro* cell culture systems are an indispensable platform for investigating cellular mechanisms of the adverse health effects of particle inhalation on the lung. Proinflammatory cytokine responses are considered a key cellular event in particles-induced diseases which mainly target pulmonary epithelial cells and macrophages. In exploring the contribution of coal particle chemistry (Song et al., 2022) to the variable risk of developing coal workers ‘ pneumoconiosis (CWP), an incurable lung disease linked to coal dust inhalation, we became concerned about protein production data which we were quantifying by enzyme-linked immunosorbent assay (ELISA). While we found a variable cytotoxic effect, this was not reflected in our cytokine assays. Upon reviewing the literature, it became apparent that there was some precedent to this observation with some studies raising concerns that the particles may non-specifically bind secreted cytokines, possibly confounding the results of bioassays such as ELISAs (Kocbach et al., 2008, Grytting et al., 2021). Thus, we hypothesised that coal particles variably absorbed extracellular cytokines, leading to underestimation of cytokine response in *in vitro* experiments.

To test our hypothesis, we first mixed pro-inflammatory cytokines IL-1β, IL-6, IL-8 and TNF-α with 200 µg/mL of coal particles in RPMI-1640 growth medium with 10 μM Phorbol 12-myristate 13-acetate and 10% fetal bovine serum (FBS) or in Ham ‘s F-12K medium supplemented with 10% FBS and 1% glutamine. The cytokine solutions were prepared from recombinant cytokine standards obtained from ELISA kits (Colorimetric assay RDSDY201 for IL-1β, RDSDY406 for IL-6, RDSDY208 for IL-8, RDSDY210 for TNF-α from R&D Systems; Fluorescent assay, ab229402 for IL-8 from Abcam). We spiked the cytokines (the second high and the last standards in the kits) into cell-free medium with high (125 ng/mL for IL-1β, 300 ng/mL for IL-6, 1000 ng/mL for IL-8 and 500 ng/mL for TNF-α) and low (8 ng/mL for IL-1β, 19 ng/mL for IL-6, 63 ng/mL for IL-8 and 31 ng/mL for TNF-α) concentrations. Five coal particle preparations were selected with homogeneous particle sizes (∼ 0.11 µm) but differing physico-chemical composition (Song et al., 2022). Mixtures of cytokines and particles were incubated for 24 h at 37°C in humidified 5% CO_2_, with a total volume of 500 µL. After incubation, the mixture was analysed for cytokine concentration with or without centrifugation (12,000 g × 5 minutes). A cytotoxicity assay (lactate dehydrogenase assay, G1780 from Promega) was also performed in a cell-free model as described for cytokine analysis.

The cytokine binding experiment in the cell-free system (Figure 1) showed that particles significantly absorbed the cytokines in the media in a dose dependent manner. Amongst the investigated cytokines, the reduction in IL-8 and TNF-α was particularly prominent. Therefore, these data support our hypothesis that coal particles are capable of binding to cytokines, which is likely to result in underestimation of their inflammatory potential. However, the binding affinity differed between coal samples and cytokines likely due to variability in the physico-chemical properties. For example, using a high dose of IL-8, the absorption varied from 14 to 69% between the 5 coal samples tested.

**Figure 1.**
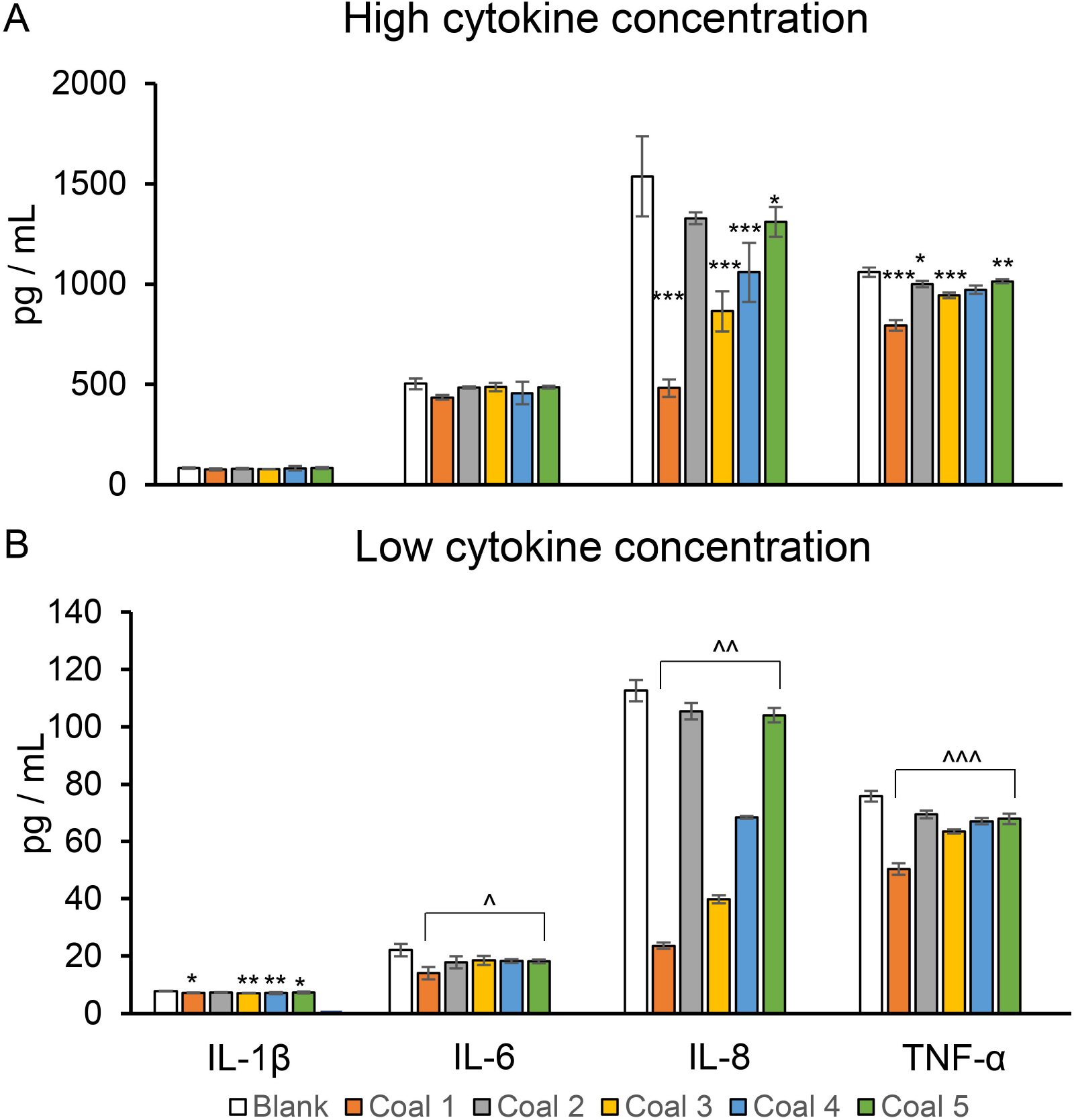
Cytokine binding experiment in a cell-free system: Inflammatory cytokines (IL-1β, IL-6, IL-8 and TNF-α) in either high (A) or low (B) concentration were co-incubated with five coal particles as well as the vehicle control (Blank) in THP-1 cell growth medium (RPMI-1640). The experiments were performed in triplicate. Difference was compared amongst the experimental groups using one-way ANOVA followed by post hoc least significant difference tests. Values are mean (SD). ^*^ *p* < 0.05, ^**^ *p* < 0.01, ^***^ *p* < 0.001 compared to the control group; ^^^ *p* < 0.05, ^^^^ *p* < 0.01, ^^^^^ *p* < 0.001, all covered groups compared to the control group.

Several investigators have identified a similar binding effect of particles on cytokines. For example, Kocbach et al. reported up to 85% binding of cytokines (TNF-α, IL-1β, IL-6, IL-8) by carbonaceous particles, including particles from outdoor sources, ultrafine carbon black and diesel but not by mineral particles (quartz) (Kocbach et al., 2008). They further found that addition of serum proteins could partly or completely reduce the cytokine binding to particles (Kocbach et al., 2008). It is worth noting that our experiment included 10% FBS and yet cytokine absorption was still very high (e.g. 69% IL-8 binding to Coal 1), indicating that this strategy may not totally resolve the problem. Grytting et al. showed that a range of respirable stone particles (400 μg/mL for 24 hours exposure) absorbed CXCL8 and IL-1β, which was of greater concern in DMEM medium, used with HBEC3-KT cells, but negligible in RPMI medium, used with THP-1 macrophages (Grytting et al., 2021). Thus, multiple factors could influence binding including particle composition, the specific cytokine (protein characteristic) and the culture conditions.

Subsequently, we tested the effect of different processing conditions / strategies on adsorption artefacts using these coal samples including the cell culture media, centrifugation, different detection methods (colorimetric and fluorescent), all of which have been reported to influence *in vitro* assay analysis (Herseth et al., 2013, Forest et al., 2015, Grytting et al., 2021). As shown in Figure 2 A-B, the absorption issue persisted in both RPMI medium and was not affected by the centrifuge protocol. It has been postulated that the interaction between particles and biological molecules interferes with absorbance measurements, confounding the results of the assay (Forest et al., 2015). However, our data showed that such interference was not improved by using a fluorescent detection method (Figure 2C). Of note, the cytotoxicity assay (LDH assay) was not affected by presence of particles (Figure 2D), which is measured using light transmission/absorbance. Thus, it is possible that the confounding of these immunoassays is likely due to blocking of antigenic regions in cytokines by the coal particles.

**Figure 2.**
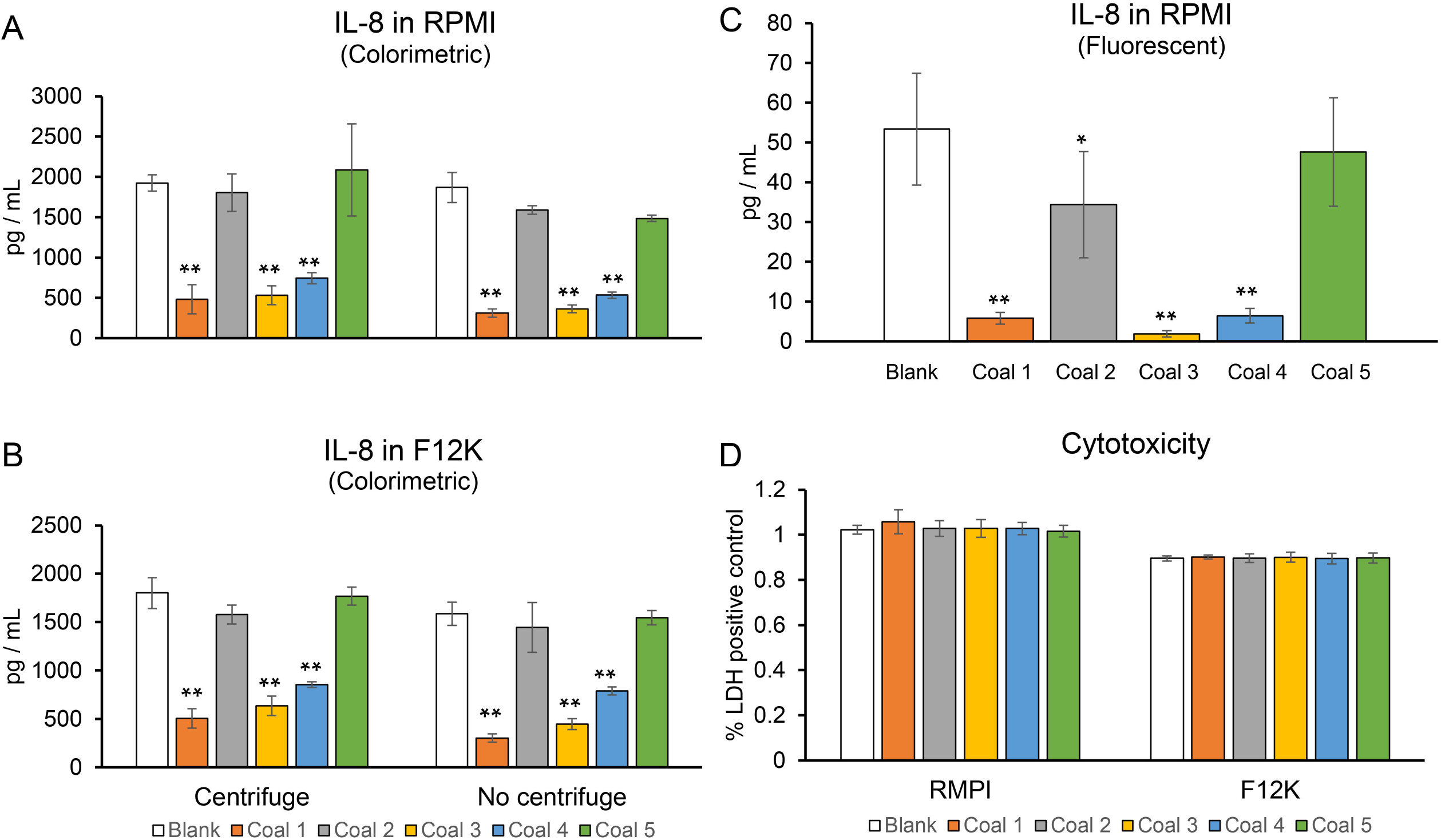
Effects of different conditions on quantification of *in vitro* cell response: IL-8 (1000 ng/mL) were co-incubated with five coal particles and the vehicle control (Blank) in THP-1 cell growth medium (RPMI-1640) and A549 cell medium (F12K), followed by either centrifuging or no centrifuging. IL-8 was quantified using Colorimetric ELISA assays (n = 3 per group; A-B). In the meantime, IL-8 quantification was performed in RPMI-1640 without centrifuging step using Fluorescent ELISA assays (n = 4 per group; C). Lactate dehydrogenase assay (LDH) was also performed in cell-free model (n = 3 per group; D). Difference was compared amongst the experimental groups using two-way ANOVA (except of fluorescent IL-8 study using one-way ANOVA) followed by post hoc least significant difference tests. ^*^ *p* < 0.05, ^**^ *p* < 0.01 compared to the control group.

In conclusion, to date, no effective strategy is available to solve this problem. We agree with previous reports that cytokine binding experiments should be performed to implement appropriate correction factors in order to calculate the accurate production of secreted cytokines in the supernatant of cell culture experiments (Kocbach et al., 2008, Herseth et al., 2013). This is an underappreciated issue in many published studies on the comparative potency of particles from different sources.

## Notes

### Competing Interest Statement

The authors have declared no competing interest.

